# Decoding depression during subcallosal cingulate deep brain stimulation

**DOI:** 10.1101/2022.07.27.501778

**Authors:** Vineet R. Tiruvadi, Ashan Veerakumar, Otis Smart, Andrea Crowell, Patricio Riva-Posse, Robert E. Gross, Cameron C. McIntyre, Christopher Rozell, Robert Butera, Helen D. Mayberg

## Abstract

Deep brain stimulation of subcallosal cingulate white matter (SCCwm-DBS) alleviates symptoms of treatment resistant depression (TRD) over months of therapy. Readouts of depression symptom severity derived from neural recordings are needed for more systematic study and improvement of the therapy. In this study, we measured local field potentials (LFP) multiple times a day alongside seven months of therapy using the Activa PC+S™ in six patients treated with SCCwm-DBS. We found significant changes in oscillatory power between early and late therapy after accounting for stimulation-related distortions, particularly within the *β* band. We then used a decoder strategy to identify oscillatory activity that tracked with depression measurements over seven months, with asymmetric *δ* and *β* oscillations contributing to a statistically significant prediction of 10% of the measured depression signal. Simulating its use in clinical decision-making, we demonstrated that the DR-SCC yield clinically meaningful information that can augment other measures of depression state. Ultimately, this DR-SCC provides a data-driven first-step towards objectively tracking chronic recovery after antidepressant DBS implantation and developing adaptive DBS strategies in the presence of active stimulation.

## 1 Introduction

Major depressive disorder (MDD) is a severe mood disturbance characterized by persistent sadness, suicidal thoughts, motor and cognitive deficits, and sleep disturbances [1, 2]. Deep brain stimulation (DBS) of subcallosal cingulate white matter (SCCwm) has been shown to be effective in alleviating treatment resistant depression (TRD) [3, 4, 5, 6, 7] However, clinical trials of antidepressant DBS have been limited in capturing efficacy, likely as a combination of unclear targeting and imperfect assessment of mood state[8, 9]. One major challenge is the reliance on patient self-report through clinical scales, a questionnaire administered to patients from trained clinical staff, as they recover along different trajectories reflecting their individual circumstances [10]. Objective, physiologically derived signals are needed to more systematically assess and improve antidepressant DBS.

MDD is a network disorder involving distributed brain regions connected directly and indirectly by white matter tracts [11, 12, 13]. The subcallosal cingulate cortex (SCC) is a particularly important node in the MDD brain network and has been shown to be metabolically correlated with mood state and antidepressant response[14, 15, 16]. This suggests an electrophysiologic correlate, potentially in the oscillatory activity of macroscopically recorded field potentials [15, 11, 17]. Early investigation of these field potentials in intraoperative [18] and post-operative [19] settings have yielded promising biomarkers, but chronic and dense recordings are needed to better capture the characteristic time constants of depression recovery. Neural recordings may serve to better reflect these behaviors, but this has been challenging due to the slow timecourse of depression recovery and the inability to measure over the same time frames probed by clinical scales and decisions [18, 19].

Physiologic signals from the brain networks underlying depression are needed [20], and recent DBS advances enable low-profile recording from around the stimulation target alongside active DBS therapy [21, 22, 23]. *Differential* LFP recording devices capable of recording alongside active therapy provide a powerful window alongside therapy, but require great care to remove stimulation-related distortions[22, 24]. Additionally, small and heterogeneous patient cohorts typical of antidepressant DBS studies require supervised machine learning to identify links of interest in the presence of unavoidable, confounding correlations. Signals likely involve several interrelated oscillations in a depression-related region, requiring multidimensional approaches to identify meaningful disease *readout*.

In this study we developed a decoding model that yielded a depression readout from chronic SCC-LFP oscillations (DR-SCC). Chronic SCC-LFP from PC+S™ were recorded in patients treated with SCCwm-DBS [25]. First, we demonstrated oscillatory changes are measured in patients before identifying personalized decoding models that achieve statistically significant performance. We then identified a parsimonious depression readout from SCC (DR-SCC) that achieved statistically and clinically significant decoding of depression state over seven months of therapy.

## 2 Methods

### 2.1 Clinical Protocol

#### Patient Recruitment

Six patients with TRD were consecutively enrolled into a study of SCC-DBS for TRD safety and efficacy (ClinicalTrials.gov Identifier NCT01984710) between 2013 and 2018 (Table 1). All patients provided written informed content to participate in the study and to have ambulatory SCC-LFP recordings sampled throughout the day. Inclusion and exclusion criteria are identical to those previously published [4, 7, 25]. The protocol was approved by the Emory University Institutional Review Board and the US Food and Drug Administration under a physician-sponsored Investigational Device Exemption (IDE G130107). The study is actively monitored by the Emory University Department of Psychiatry and Behavioral Sciences Data and Safety Monitoring Board. All patients participated in a study involving multiple LFP recordings per day over the course of at least seven months post DBS implantation.

**Table 1:** SCC-LFP Patient Recordings. Patient Demographics and number of recordings available from each patients over the first seven months post-implantation.

|  | Sex | Age | Baseline HDRS17 | 6mo HDRS17 | 12mo HDRS17 | Recordings |
| --- | --- | --- | --- | --- | --- | --- |
| Pt 1 | F | 50 | 23.5 | 6 | Y | 1287 |
| Pt 2 | F | 48 | 20.25 | 7 | Y | 1433 |
| Pt 3 | F | 70 | 23.25 | 15 | Y | 801 |
| Pt 4 | M | 64 | 19.25 | 6 | Y | 794 |
| Pt 5 | F | 62 | 22.75 | 7 | Y | 714 |
| Pt 6 | M | 57 | 20.5 | 10 | Y | 761 |

**Table 2:** Naive Oscillatory Changes over Six Months of Therapy. Number of patients that exhibit increases and decreases at each oscillatory feature. MC uncorrected (top two rows) and MC corrected (bottom two rows) show impact of stimulation artifact.

| | L-δ | L-θ | L-α | L-β | L-γ | R-δ | R-θ | R-α | R-β | R-γ |
| --- | --- | --- | --- | --- | --- | --- | --- | --- | --- | --- |
| ↑ uncorrected | 3 | 3 | 4 | 4 | 4 | 2 | 3 | 3 | 3 | 4 |
| ↓ uncorrected | 0 | 2 | 1 | 1 | 0 | 1 | 2 | 2 | 2 | 0 |
| ↑ corrected | 4 | 0 | 1 | 1 | 1 | 4 | 1 | 2 | 2 | 0 |
| ↓ corrected | 1 | 1 | 2 | 4 | 4 | 0 | 2 | 3 | 4 | 4 |

#### Implantation and Management

Patients were stereotactically implanted with DBS leads (Medtronic DBS3387) bilaterally in SCC [7]. Individualized SCCwm targets were determined through both MRI and tractography in a protocol previous described (Figure 1a) [7]. One of the middle two DBS electrodes was placed at the SCCwm target to allow for differential LFP recordings. Therapeutic DBS was initiated one month after implantation to allow for post-operative healing and maintained chronically for 24 weeks. Stimulation at the therapeutic electrode was initiated at 3.5 V and adjusted by study psychiatrists after clinical assessment. All other parameters were fixed: bilateral stimulation at 130 Hz, 90 µs pulsewidth, unipolar stimulation with IPG-cathode.

**Figure 1:**
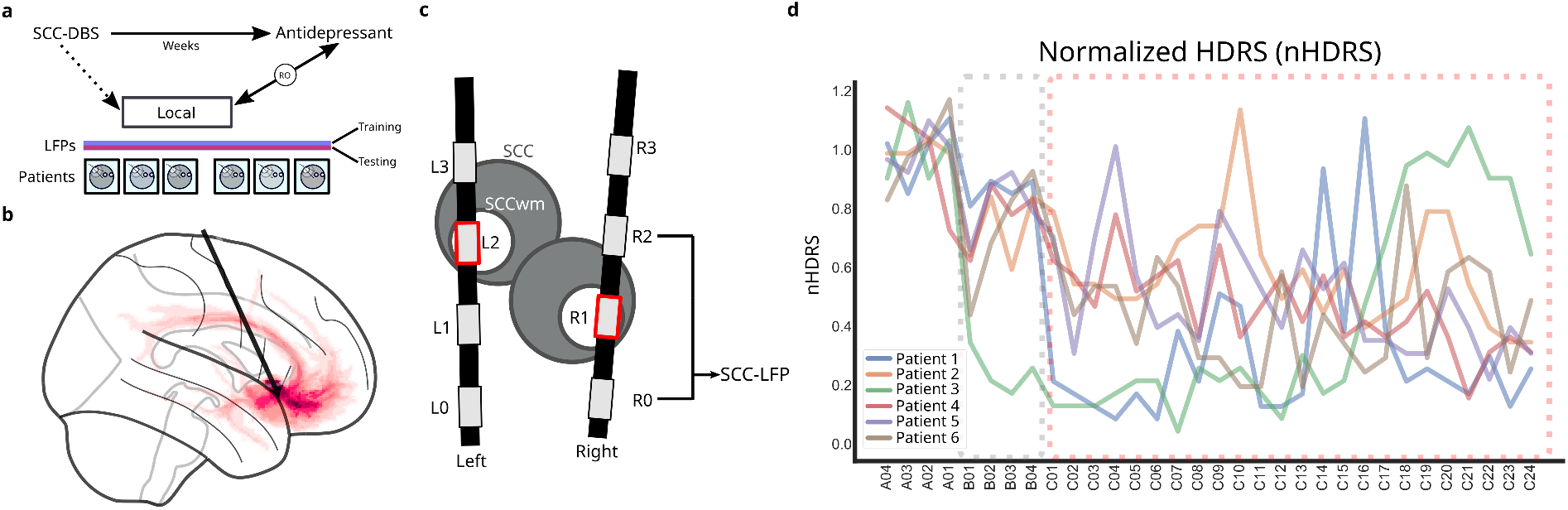
Overview of SCCwm-DBS and Activa PC+S™ recordings. **a**, SCCwm-DBS alleviates depression over weeks and months of therapy. Recordings taken locally at the SCCwm reflects oscillations at bilateral SCC, providing data to identify a *depression readout*. LFPs are recorded over seven months in six patients, split into training and testing sets, and analysed with linear regression approaches. **b**, DBS is targeted at tractography guided SCCwm, bilaterally. **c**, Bilateral leads allow for two differential LFP recording channels using electrodes around the chronic stimulation electrodes (red).

**Figure 2:**
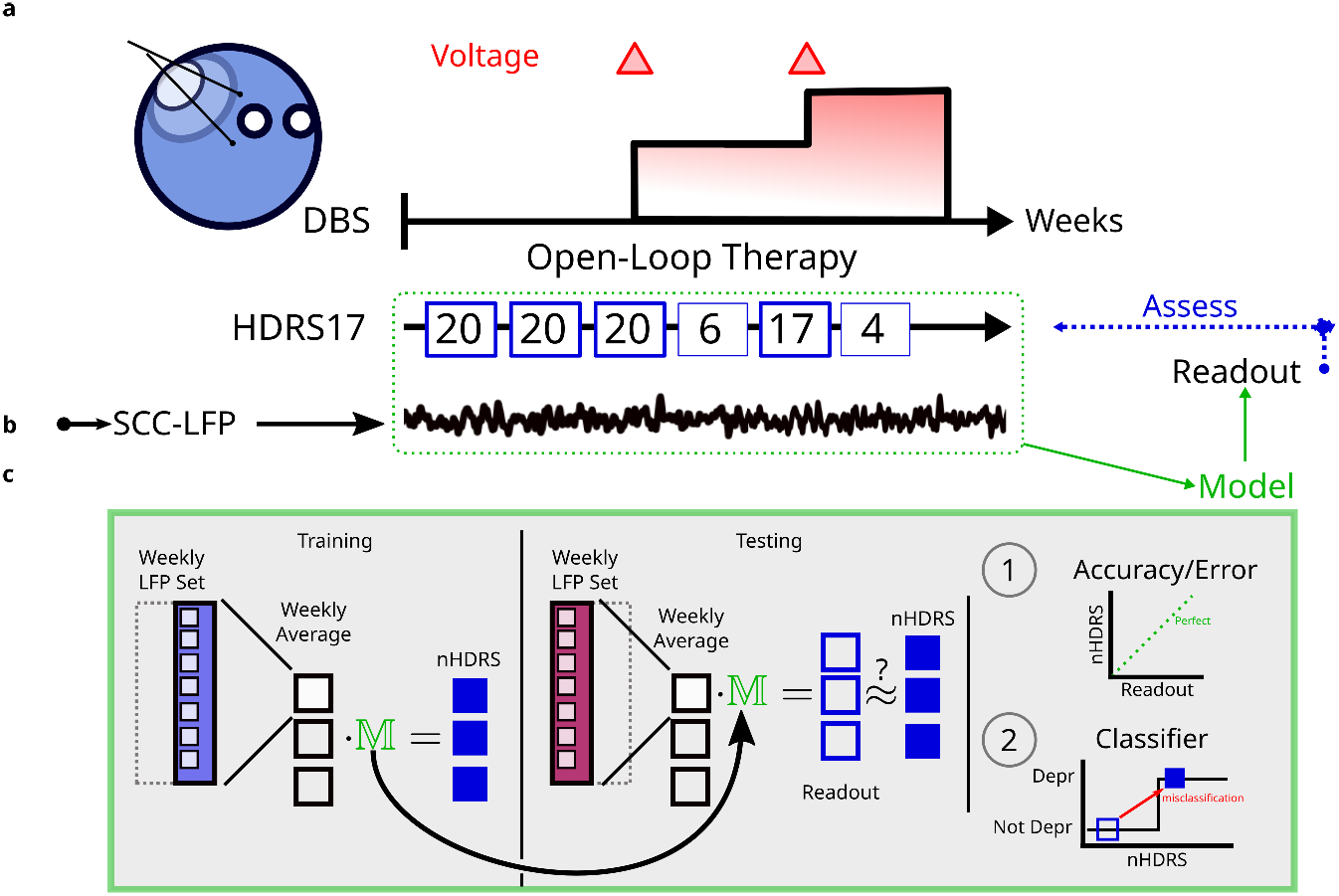
Methods Overview. **a**, Therapy settings are determined solely through clinical assessment alone. **b**, *∂*LFP recordings are measured multiple times a day in parallel. **c**, Recordings are used to train and test a regression model. Readout correlation with nHDRS is assessed. Additionally, simulated clinical decision-making using the readout is assessed against clinical decision making.

#### Weekly Depression Scale

The HDRS17 is a series of 17 questions that assesses the severity of several depression-specific and non-specific symptoms over the previous seven days. An HDRS17 was administered weekly by trained clinical staff [26]. For each patient, treatment response was defined as a 6-month HDRS17 below half of the average pre-implantation HDRS17. The HDRS17 was used to assess treatment response at the study endpoint, though it was not used alone to make clinical decisions. To enable comparison between patients, each patient’s empirical HDRS17 is divided by the four week pre-surgical average, yielding a normalized HDRS (nHDRS). After normalization, an nHDRS below 0.5 at six-months corresponds to treatment response. n=5/6 patients were responders at six months and all six patients were responders at twelve months.

### 2.2 Neural Recordings

#### Local Field Potentials

Local field potentials (LFPs) were recorded with the Activa PC+S™ (Medtronic PLC., MN, USA), a dual-channel recording device that measures *differential* LFP [27, 24]. Two channels were recorded at 422 Hz, one each in left and right SCC (Figure 1c). dLFP channels are formed by the two electrodes directly adjacent to the stimulation electrode, both 1.5 mm away, enabling common-mode rejection of stimulation artifacts [22]. Hardware bandpass filters from 1 Hz to 100 Hz and amplifier gains were set per-channel as the highest of four options: 200,400,1000,4000 without overt saturation, as determined by Medtronic engineers. 15 s to 20 s recordings are measured approximately every six hours and stored on-device, with intervals tuned to maximize number of recordings before the next scheduled visit without overfilling onboard memory. LFPs are measured both without stimulation (the four weeks post implantation) and with stimulation (twenty-four weeks after therapy initiation).

#### Preprocessing

The final 10 s of each recording is used for all analyses to avoid an amplifier settling period in the first 2 s to 5 s. Daytime recordings collected between 8am and 8pm were isolated for analysis to ignore circadian and focus on period of time most likely to reflect the symptoms probed by the HDRS17 [28].

To remove distortions related to mismatch compression, an extra preprocessing pipeline was applied in the frequency-domain representation of all recordings, unless otherwise specified. This pipeline removes broad-spectrum features, narrows frequency bands for oscillations, and measures the level of gain compression driven by stimulation-related distortions in the presence of heterogeneous tissue between the two *∂*LFP recording electrodes [24].

#### Weekly States

Ultimately, six patients had X weeks of brain (LFP) and behavior (nHDRS) measurements. All recordings taken within a given week are associated with the immediate next nHDRS sampled. This is justified by the nHDRS specifically assessing the previous seven days of symptom. The total number of LFP recordings o=5790 are pooled across all patients, with each recording linked to the patient’s nHDRS in the immediate next clinical assessment.

#### Oscillatory State Calculation

Oscillatory state is computed for all recordings as the power in each oscillation Oscillatory states are calculated from each recording as the power in oscillatory bands across both SCC-LFP channels. Recordings are transformed into power spectral densities (PSDs) using the Welch algorithm with Blackman-Harris window of 2 s, 0% overlap and 2^10^ = 1024 FFT bins. The PSD is then preprocessed to remove mismatch compression distortions [22, 24] arising from chronic impedance variability. Average (median) oscillatory power is calculated within five modified band windows: *δ* (1-4Hz), *θ* (4-8Hz), *α* (8-14Hz), *β*^*\**^ (14-20Hz), *γ*^1^ (35-50Hz). The frequency ranges for higher oscillations are adjusted to avoid stimulation-related device artifacts and the device noise-floor. The final brain state is the full feature vector of all 10 bilateral SCC features.

### 2.3 Depression Decoder

#### Linear Regression

The decoding model takes the bilateral SCC oscillatory power as an input to generate a single scalar value as output, the *depression readout*. Elastic net regularized linear regression is used to train a linear decoder to identify first-order links between SCC and depression recovery in a way robust to collinearities amongst oscillations and encouraging of sparsity[29]. The DR-SCC decoding model is trained on weekly averaged oscillatory states in the training set across all patients (Figure 4a). Patient-level cross-validation (leave-three-out) yields 20 sets of model coefficients, distributions depicted in violin plots (Figure 4). Feature-wise mean is taken as the final DR-SCC model (Figure 4a, blue line).

#### Implementation

ENR is implemented through the ElasticNet class in scikit-learn [30] and ENR hyperparameters are chosen to optimize between model fit *R*^2^ and the number of non-zero coefficients in the model (See Supplementary 6.4; Figure S10).

#### Training/Testing Split

Daytime recordings, and their oscillatory states, are then randomly split into non-overlapping training (60%) and testing (40%) sets. The SCC states for all recordings are calculated and then binned into their appropriate clinical week. The feature-wise median is calculated for all recordings taken within the same patient and clinical week, yielding a single average oscillatory state for each week in the set.

#### Hyperparameters and Cross-Validation

In training the depression decoding model, hyperparameter optimization and patient-level cross-validation is implemented. CV is used to train on all combination of three patients, yielding a total of 20 trained models and a distribution of coefficient values. ENR hyperparameters are chosen systematically to maximize *R*^2^ fit while minimizing the number of variables included in the model (Supplementary Figure 11). The final decoding model is calculated as the coefficient-wise mean across CV folds, yielding the parsimonious decoding model and DR-SCC readout.

#### Statistical assessment

Assessment of the DR-SCC is done in a held-out testing set of individual recordings over the same time period, in the same patients. The testing set is randomly subsampled over 100 iterations, each iteration including a random subset of 80% of the testing set recordings [31, 32]. The model was assessed by randomly subsampling 80% of the testing set recordings, calculating weekly average oscillatory states, and assessing the DR-SCC against the empirical nHDRS using an actual-vs-predicted plot [33]. The prediction’s *R*^2^ then reflects the percent of the actual explained by the readout, and statistical significance is assessed as *p <* 0.05. An additional analysis is performed comparing predicted-vs-actual using surrogate nulls by permuting the recording set against a fixed nHDRS set (See Supplementary 11).

### 2.4 Clinical Assessment

#### Clinical Significance

To assess the clinical significance of the readout, we simulated its use in clinical decision making using simplified control strategies. We simulated two clinical tasks: response classification and DBS amplitude increase prediction.

#### Response Classification

Therapeutic response is defined as a nHDRS *<* 0.5 at the six month time point. A *receiver-operator curve* analysis is performed on the DR-SCC to assess its congruence with the nHDRS determined treatment response at every clinical week. The area-under the ROC (AU-ROC) was calculated over 100 trials of randomly subsampled observations in the testing set to yield a distribution of performances, with the mean performance of the distribution assessed against pure chance (AU-ROC = 0.5).

#### DBS Amplitude Increases

DBS amplitude is increased by clinicians after a clinical assessment, a complete assessment of the patient’s depression state. This serves as a useful surrogate label for the depression state of the patient: no stimulation increase is interpreted as not depression.

#### Simulated Control Strategies

A control strategy consists of a measure of depression paired with a control policy. We assessed a set of measures of depression that included the putative depression readout, along with the nHDRS, a null, and an oracle with noise. We implemented a single threshold control policy: if a readout was above threshold then a positive event is predicted. The predicted positive events are then compared to the empirical positive events using a precision-recall curve (PRC) analysis, due to class imbalances in the stimulation increases, occurring only 8/128 observed weeks. The PRC analysis varies the threshold between 0 and 1, calculating the precision () and recall () in predicting the presence of an amplitude increase at each week. The average precision is assessed as well as the PRC across thresholds, and compared between readouts from the readout set.

### 2.5 Analysis Code

Preprocessing and analysis are implemented in Python using standard (`numpy, scipy`) libraries [30, 34]. Linear regression training, cross-validation, and assessment is implemented through the scikit-learn library [30]. All analysis code is available open-source at github.com/virati/SCC_readout. Raw and processed data available upon request.

## 3 Results

### 3.1 Oscillatory Changes

#### Power Spectral Density Changes

Changes in the power spectral density (PSD) of weekly averaged recordings are compared between the first (C01) and twenty fourth (C24) weeks of therapy (Figure 3). Several artifacts are evident in the PSDs, including an 32 Hz mismatch compression related artifact, and a device-related 25 Hz artifact (Figure 3a,b). Right-SCC PSDs exhibited significantly more variability than Left-SCC PSDs (Figure 3b). The peak found at 25 Hz was found during *in vitro* recording as a voltage-sensitive artifact [24]. This peak is included in traditional *β* oscillation definitions (15 Hz to 30 Hz) but is ignored as a part of mismatch compression corrections *β*^*\**^ (15 Hz to 20 Hz) [24].

**Figure 3:**
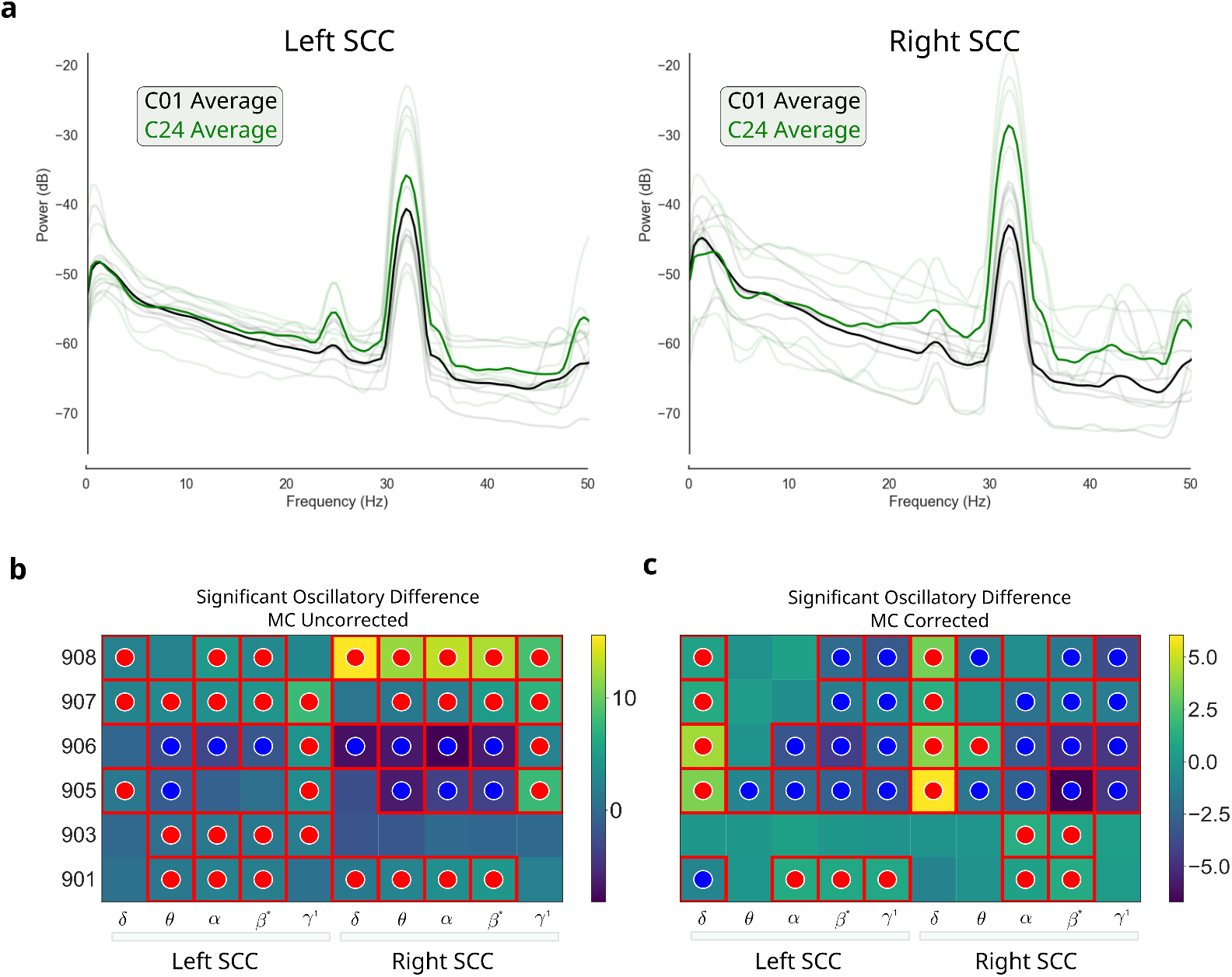
Oscillatory Power Changes over Therapy. **a**, Weekly average power spectral densities (PSD) between the early (black solid) and late (green solid) therapy courses, for all patients (translucent lines). **b**, Significant oscillatory band changes over the time frame, without corrections for mismatch compression. **c**, Significant oscillatory band changes over the time frame, with corrections for mismatch compression.

#### Uncorrected Oscillations

Oscillatory power changes between C01 and C24 calculated without mismatch compression corrections showed consistent changes in *θ, α*, and *β* oscillations bilaterally in at least 5/6 patients (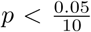 ; Figure 3b).

#### Corrected Oscillations

Oscillatory power changes between C01 and C24 after correcting for mismatch compression showed consistent *L − δ, L − β*^*\**^, *L − γ, R − α, R − β*^*\**^ all exhibit changes in at least 5/6 patients (Figure 3c). Right *β\** changes in all patients, with predominant decrease (Figure 3c).

### 3.2 Depression Readout from Subcallosal Cingulate Cortex (DR-SCC)

#### Readout Performance and Statistics

A single, parsimonious decoder was trained from weekly averaged SCC oscillatory states (Figure 4). The predictive performance of the DR-SCC, explaining approximately 10% of the nHDRS variance in a held out testing set, is statistically significant (Figure 4; *p <* 0.05). The *R*^2^ is approximately .11, with a testing-set distribution ranging from -0.05 to 0.30, indicating approximately 11% of the depression trajectory is captured by the full DR-SCC pipeline. A single sample trial is depicted in Figure 4c, with associated *R*^2^ and Pearson scores. The DR-SCC statistical significance was determined through a distribution of the readout *R*^2^ (Figure 4d). This procedure was run 100 times to yield a distribution of *R*^2^ with mean 0.11.

**Figure 4:**
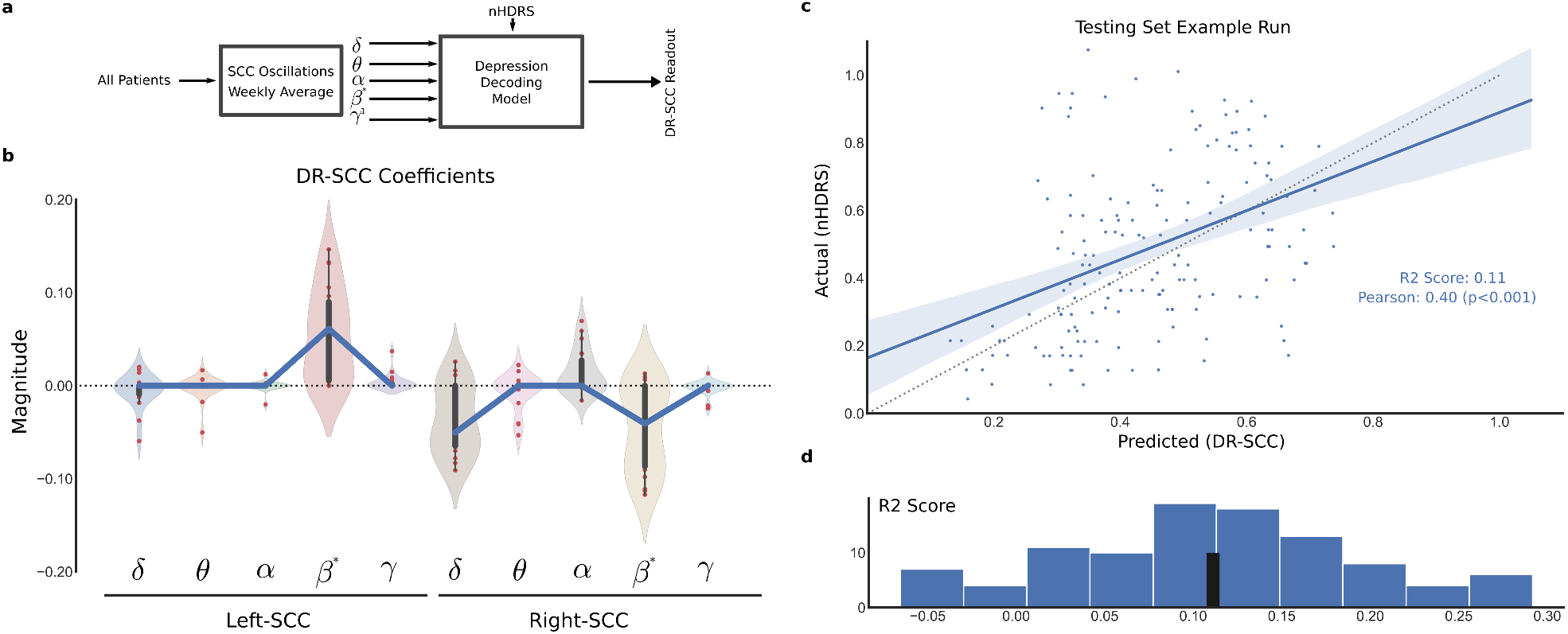
DR-SCC Decoding and Readout. **a**, A single decoding model is trained on testing set observations from all patients. **b**, Distribution of coefficients from all CV folds. Coefficients are averaged (blue line) to yield the final decoding model. **c**, DR-SCC explains approximately 10% of the empirical depression measure, demonstrated in a single testing set assessment. **d**, 100 runs of subsampled testing sets yields a distribution of performances.

#### Decoder Coefficients

The DR-SCC identified three major oscillations of interest: left-*β*^*\**^, right-*β*^*\**^, and right-*δ* 4b). Asymmetry between left and right is observe, with left-*β*^*\**^ having positive loadings indicating positive correlation with depression severity, and right-*β*^*\**^ and right-*δ* having negative loadings indicating negative correlation with depression severity. Regularization path analysis identifies the order that other oscillatory features drop out of the final model (Figure S10). A secondary analysis rules out stimulation artifact as a significant contributor to the DR-SCC (Figure S8).

### 3.3 Clinical Significance of DR-SCC

#### Binary Response

We tested a binary classification of patient response and compare it to the empirical nHDRS (Figure5a). An ROC curve is calculated over 100 trials of 80% subsampled weeks from the testing set, yielding a distribution of ROC curves (>0.5; Figure5b) and AUC measures (Figure5c). The average AUC achieved across iterations was approximately 0.7, demonstrating a statistically significant ability to predict the response.

**Figure 5:**
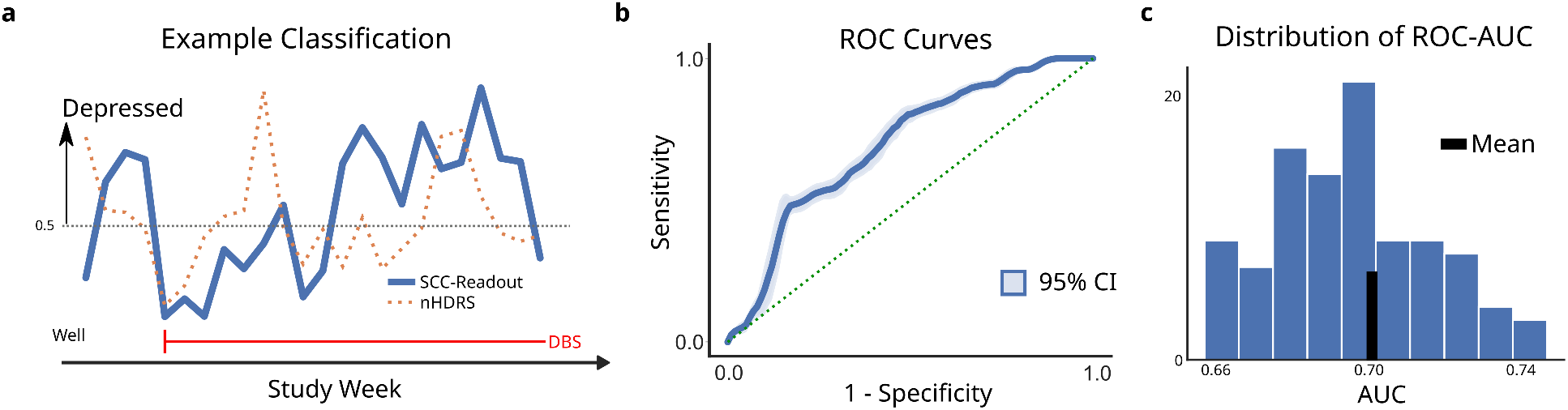
Response Classification. **a**, Illustration of a single prediction trial over 28 weeks. nHDRS scores (orangedotted) are used to label weeks as either ‘depressed’ or ‘not depressed’. The agreement of the readout (blue-solid) is then assessed through an ROC curve. **b**, ROC curves are calculated for 100 validation trials with randomly sampled subsets of the testing set. Mean ROC curve (blue line) and its 68% confidence interval (blue shade). All ROC curves were significantly above chance (green-dotted). **c**, The distribution of AU-ROCs over 100 trials ranged between 0.66 and 0.74, with median of 0.70.

#### Simulated Adjustments

To see how the DR-SCC would perform if we had made decisions to increase stimulation amplitude based off of it, we simulated its use in a simple control policy (Figure 6). Here, the policy is that a suprathreshold DR-SCC warrants a stimulation increase and the threshold is swept across all values to calculate a precision-recall curve (PRC). Over a total of 128 observations, a total of eight stimulation increases were administered across the cohort (Figure 6a). A total of four policies (metric + strategy) were assessed: nHDRS alone, DR-SCC alone, Null, and Oracle with noise (Figure 6b). An oracle with a fixed noise level is used as a tractable best case scenario. The DR-SCC, in isolation, performed worse than all other estimate of the depression state, including the null and nHDRS.

**Figure 6:**
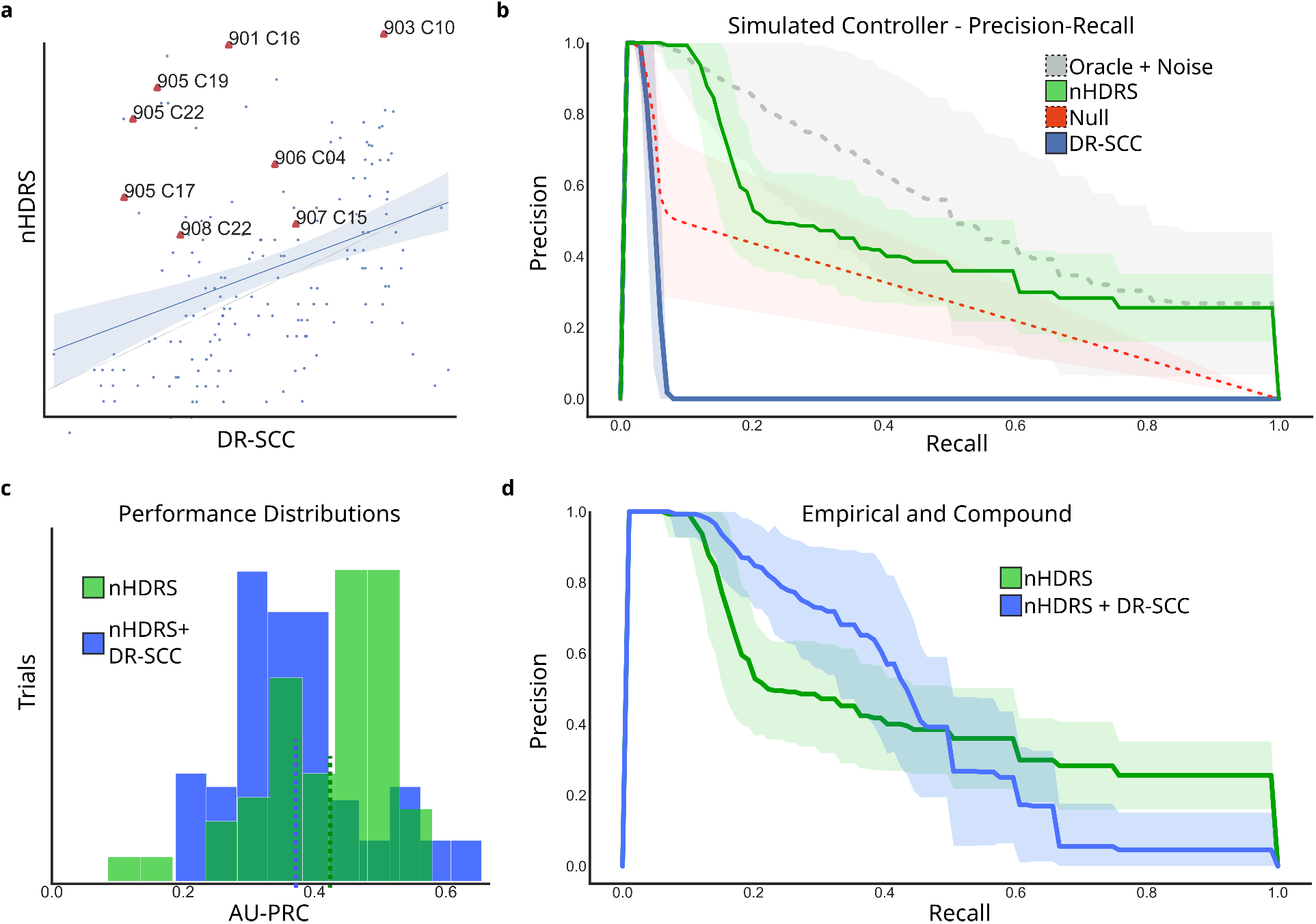
Response Classification. **a**, Illustration of a single prediction trial over 28 weeks. nHDRS scores (orange-dotted) are used to label weeks as either ‘depressed’ or ‘not depressed’. The agreement of the readout (blue-solid) is then assessed through an ROC curve. **b**, ROC curves are calculated for 100 validation trials with randomly sampled subsets of the testing set. Mean ROC curve (blue line) and its 68% confidence interval (blue shade). All ROC curves were significantly above chance (green-dotted). **c**, The distribution of AU-ROCs over 100 trials ranged between 0.66 and 0.74, with median of 0.70.

#### Augmented Decisions

The role of the DR-SCC is assessed as an adjuvant to the nHDRS, where suprathreshold values of both are used to predict a stimulation parameter increase (Figure 5f,g). The nHDRS alone yields an average PR-AUC of 0.42, while the nHDRS w/ DR-SCC yields a PR-AUC of 0.38 (Figure 6c), with the adjuvant DR-SCC approach yielding some trials with PR-AUC higher than 0.6. When inspecting the PR curves directly, a range of thresholds is evident where the precision of the adjuvant DR-SCC is higher for a fixed precision than the nHDRS alone (Figure 6d).

## 4 Discussion

In this study, we used a novel set of chronic SCC recordings taken alongside antidepressant SCCwm-DBS to identify oscillatory power changes reflecting recovery from depression. Six patients were implanted with Activa PC+S™ IPGs, with 5/6 treatment responders at six months (Table 1). Alongisde clinical management of DBS parameters, we recorded SCC-LFP multiple times a day in order to link oscillatory changes to weekly changes in HDRS17 clinical scores. In this report, we analysed recordings taken over the first seven months post-implantation, with a one-month post-implantation period without active stimulation. After accounting for mismatch compression distortions present in recordings taken with active stimulation we observed several convergent lines of evidence that coordinated SCC oscillatory activity predicts a significant part of the depression recovery trajectory.

### 4.1 Oscillations change over six months of therapy

While SCC metabolism has been strongly linked to depression state and treatment response [14, 15], the presence of electrophysiologic correlates of depression in SCC recordings remains unclear. Early investigations limited to intraoperative [18, 35] and semi-chronic [19, 36] recordings have identified oscillatory features that track with specific symptoms of depression, but a complete readout requires recordings over the weeks and months of mood recovery. To address this, we first identified oscillatory changes occurring over two critical milestones: the first week of active stimulation (C01) and the six month timepoint (C24).

SCC oscillations demonstrated significant changes across six months of therapy (Figure 3), both with and without correction for mismatch compression [24]. After adjustments, particularly in *β →β*^*\**^ to avoid a narrowband peak, we found consistent changes in *δ* and *β*^*\**^ over the study, though significant variability across patients is evident (Figure 3). Variability in the depression recovery trajectories may account for this observation (Figure 1d) but numerous other sources of variability may be confounding a simple early-vs-late analysis. Account for large source of noise is a necessary first step in establishing reliable chronic depression readouts, and our development of a mismatch compression correction pipeline was critical to this end [37, 38, 24].

### 4.2 Decoder Strategy Identifies Depression Correlates

We used a decoder strategy to more directly link SCC oscillations to depression state by leveraging observations at all weeks along the recovery trajectory [39, 40]. To more directly link measured SCC oscillations to depression state, we took a decoder strategy using recordings from all available weeks. These predictions, while modest, achieved statistical significance. The resulting correlation suggests the SCC accounts for only part of the, consistent with MDD being a disorder arising across distributed networks. Improving on this performance for future readouts may require coverge of brain regions beyond SCC, or computational models that can infer network-wide states from limited SCC observations [41].

While only accounting for 10% of the total variance, the DR-SCC achieves a statistically significant prediction. One interpretation of this is that bilateral SCC only accounts for a subset of all the symptoms assessed by the nHDRS, so full correlation with the nHDRS using only SCC recordings may be untenable [11, 12, 13]. Improving on the DR-SCC explained variance will require either more measurements or the use of whole-brain models for inference [19].

### 4.3 DR-SCC implicates *β* and *δ*

The decoder model driving the DR-SCC identifies two oscillations of interest: *δ* and *β*^*\**^. These coefficients are robustly present across all cross-validation folds suggesting a conserved signal across patients. A stark asymmetry is observed across bilateral *β*^*\**^ coefficients. While hemispheric asymmetry has long been associated with depression trait those asymmetries have been observed in *α* along surface EEG. *β* is a central oscillation of study in other DBS indications, like Parkinson’s Disease and Essential Tremor. One explanation of these seemingly disparate priors is that the same underlying activity can manifest as *α* across larger loops involving surface cortex and higher-frequency *β* loops in deeper structures with smaller loops.

Several factors explain the 10% explained variance. First, we ignore *γ* oscillations due to technical limitations int he PC+S™ that prevent reliable high-frequency analysis. Second, we are measuring only from the SCC and depression is known to involve a larger brain network. Third, the HDRS17 measures symptoms that are both specific and non-specific to depression, and failure to explain non-specific subcomponents of the HDRS17 would be classified here as an error.

### 4.4 Noise Invariance

The use of supervised machine learning, via regularized regression, enabled reliable identification of neural oscillations that track with depression specifically instead of other clinically imposed confounds. A major strength of this analysis is the use of a regularization, which imposes both sparsity and stability on the final decoder, accounting for potential sources of noise in both the neural recording as well as the nHDRS measurement of depression [29].

As a secondary analysis and confirmation that the DR-SCC is not being driven by stimulation artifact, we explicitly included a measure of mismatch compression in the feature set and re-ran the decoder training (Figure S8) [24]. The model itself was very similar to the decoder and the performance did not change appreciably. This supports a neural driver of the DR-SCC, but further validation in next-generation devices is needed.

### 4.5 DR-SCC has some clinical utility

To assess the clinical significance of the DR-SCC, we simulated its use in two clinical tasks: assessment of treatment response, and prediction of therapeutic voltage increases (Figure 6 and Figure 5). First, had we determined the treatment response of the patients using the DR-SCC instead of the nHDRS, our decisions would have aligned significantly with the empirical clinical assessments made in the study (Figure 5a-c) and the stimulation voltage increases (Figure 5d-g) using the DR-SCC and the nHDRS. This supports the presence of a meaningful depression signal in measured oscillatory power.

Second, we assessed the DR-SCC ability to predict increases in therapeutic stimulation frequency following a clinical assessment of depression or risk of depression (Figure 5d). Comparison of various simulate controller policies demonstrates variable performance in their precision-recall curves, though interpretation can be challenging [42]. The oracle + noise served as an optimal comparison due to the inaccessibility of parts of the PRC space [42]. Compared to oracle and nHDRS performance, the DR-SCC alone performed substantially worse than null (Figure 5e), indicating its inappropriateness, alone, as a readout. When used as an augment to the nHDRS, however, the measure demonstrates some inconsistent benefits, seen as a broadening of the AU-PRC testing set distribution (Figure 5f). However, when combined with the nHDRS, the DR-SCC enables an improvement in predicting the stimulation changes, reflected in a right-ward shift of the PR curve (Figure 5g), though the average PR is reduced from the nHDRS-only policy (Figure 5f). Together, these suggest that the DR-SCC captures a meaningful depression-related signal in the SCC, but that it does not capture all the information needed to inform clinical decision-making algorithmically.

### 4.6 Limitations

First, the small number of patients limits generalizability to broader antidepressant DBS or TRD. The cross-validation analysis suggests the DR-SCC is robust within the cohort, but the training-testing split of recordings taken from the same weeks may inflace prediction accuracy. Second, the testing-set performance achieves an *R*^2^ of 0.11, reflecting approximately 11% of the depression signal being accounted for by the entire decoder. Several factors may explain this, including limited recording of the complete depression network, the use of linear regression models, and the technical limitations of the PC+S™ in the *γ* range [27, 43, 44, 45]. Efforts were taken to rule out stimulation-artifact as a driver of DR-SCC performance (Figure S8) but it remains possible that artifacts contribute to the DR-SCC. Third, limiting features to the oscillatory power was necessary due to the filtering hardware of the PC+S™ preventing reliable analysis of oscillatory phase. Fourth, the demonstration that the DR-SCC has some utility when augmenting the nHDRS leveraged a simple threshold classifier, like those being used for adaptive DBS in Parkinson’s Disease [46]. A more sophisticated analysis may be warranted to directly regression SCC oscillations against stimulation parameter changes, though larger datasets and cleaner recordings are likely necessary.

### 4.7 A first order readout

Despite the limitations, these results are a novel first demonstration of decoding depression state across chronic timescales from intracranial recordings of the SCC in the presence of active DBS. Explicit mismatch compression corrections enabled the use of hourly recordings sampled evenly across weeks, more regularly capturing symptoms probed by the HDRS17 [22, 24]. Additionally, the identification of oscillatory correlates during active stimulation may capture depression-salient brain dynamics present only during active stimulation drive, complementing recent results from our group [28, 47]. The observation of changes within the *β* range is consistent with other DBS indications [48] as well as with resting-state results from our own lab [47, 18]. The asymmetry observed aligns with asymmetries found in depression, though these typically involve frontal *α*, and with asymmetries evoked by SCCwm actions on brain networks [49]. This preliminary decoder can inform further hypothesis-driven investigation of neural dynamics tracking depression state [50].

## 5 Conclusion

We used a novel set of chronic intracranial recording taken over months of active DBS to identify an oscillatory readout of depression from subcallosal cingulate cortex oscillations. Consistent decreases were found within *β* oscillatory power bilaterally when comparing states at six months versus the start of the study. A weekly decoder identified asymmetric weights across bilateral *β* and right *δ*, with left decreases and right increases in power being associated with therapeutic response. This DR-SCC was able to explain approximately 10% of the depression measurement, achieving both statistical significance and clinical relevance. Ultimately, these results in a novel dataset and approach yield a preliminary readout from chronic SCC oscillations that can be further studied and extended to better link oscillatory activity in the brain to depression state.

## Acknowledgment

Thank you to the clinical team, particularly Sinead Quinn, and Lydia Denison. Thanks to Scott Stanslaski at Medtronic for technical assistance with the Activa PC+S™ device. Thank you to Dr. Allison Waters for assistance in experimental data collection. A final, most important thank you to the patients who were critical collaborators in this work.

## 6 Supplementary

### 6.1 Individualized Decoders

Decoders were first developed within patients to demonstrate proof-of-principle and improved utility over timepoint comparisons (Figure 7). In particular, avoiding the dichotomization of depression state and, instead, leveraging the entire HDRS measurement We developed patient-specific decoding models on individual recordings as a proof-of-principle that recordings contained depression-relevant information throughout the recovery trajectory, avoiding detrimental dichotomization [51].

**Figure 7:**
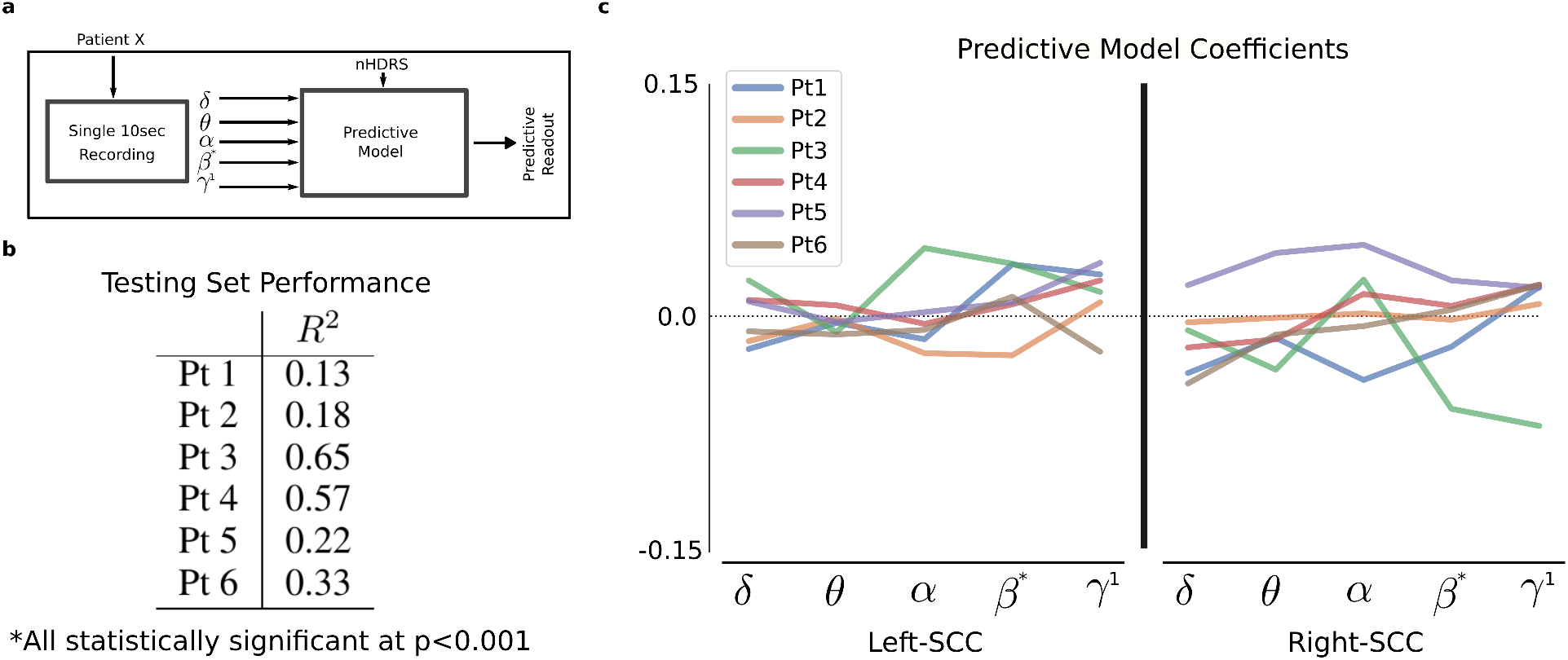
Individualized predictive model performance. **a**, Models were trained within a given patient using ordinary least squares (OLS) to yield a predictive model. **b**, This yielded a set of individualized SCC-readouts that demonstrate the highest testing set accuracy observed in this study.

#### Per-recording Prediction

Personalized decoding models were trained and assessed for ML proof-of-principle (Figure 7). Recordings were limited to individuals (Figure 7a) and used directly to train, assess, and analyse a decoding model (Figure 7b,c). Model performance, measured by *R*^2^ ranged from 13% to 65% nHDRS variance explained (Figure 7b). Statistical significance was assessed through 100 random permutations to assess performance against a simulated null (*p <* 0.005; Supplementary Figure 6.5). In all cases, the personalized decoding model was better than chance, though the percent of the variance explained was variably.

#### Decoder Coefficients

Model coefficients for all personalized decoding models were inspected to identify contributions of individual oscillations. Model coefficients for all patients exhibited significant variability, with right-SCC variability larger than left-SCC (Figure 7c). Broadly, left-SCC coefficients were smaller than right-SCC coefficients (Figure 7c). Differences in coefficient sign between patients were evident in all features, limiting parsimonious patterns.

#### Comments

Leveraging linear regression across all our oscillatory measurements, we performed linear regression between SCC oscillatory states and nHDRS17 (Figure 7) within patients at the level of individual recordings. The resulting individual models demonstrate a range of prediction performances (Figure 7a). Importantly, since recordings in both the training and testing set are taken from the same time period, the autocorrelations between observations likely inflate the performance. The right-SCC coefficients exhibited larger variability between patients than the left-SCC (Figure 7c), consistent with the larger variability in the right-SCC PSDs (Figure 3a, right panel). The range of testing set performances, combined with the asymmetry of the coefficients across patient-predictive models, strongly suggests a neural signal is being measured and that this signal can be used to achieve meaningful tracking of depression state.

### 6.2 DR-SCC robust to stimulation artifact

While steps are taken to remove features distorted by stimulation artifact, an additional analysis was performed to assess sensitivity to stimulation artifact directly. The model was retrained with a *gain compression ratio* included as a feature reflecting the level of gain compression occurring secondary to stimulation artifacts. The resulting regression model is consistent with the SCC-readout model developed in Section3.2 with a similar readout performance (Figure8). The weights on GCr features is zero

**Figure 8:**
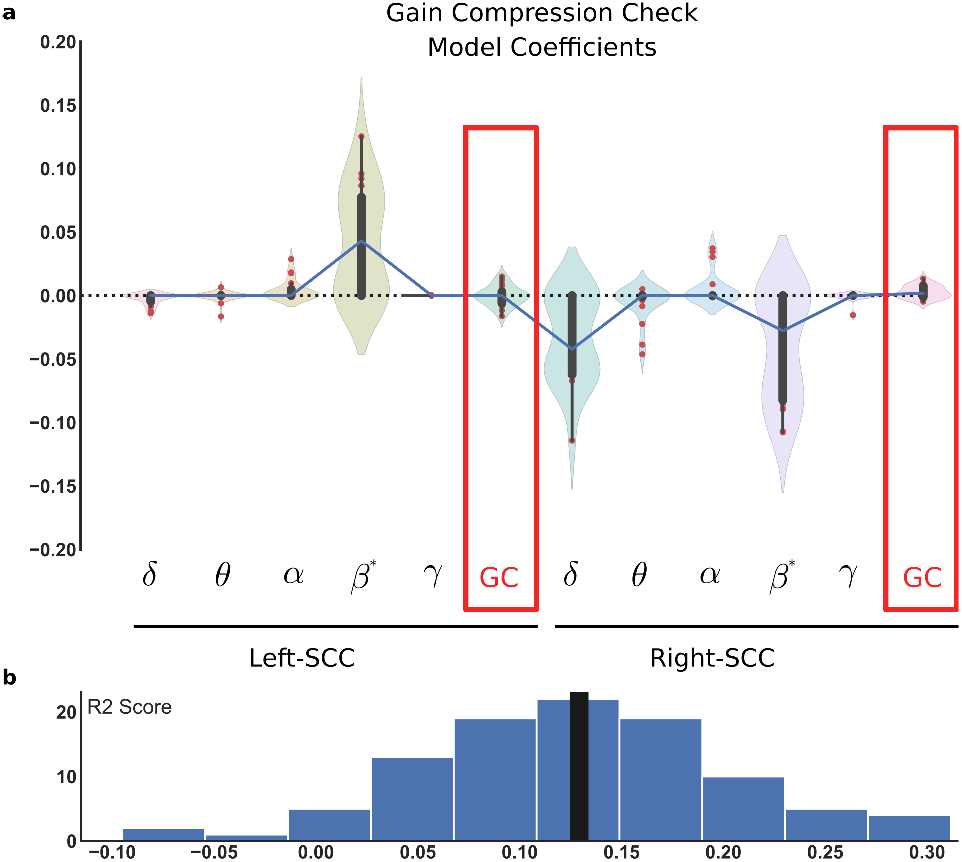
Regression with stimulation artifacts included as features. a) The entire procedure is re-run, this time including a measure of nonlinear stimulation distortion. No significant change is observed in the model parameters. b) No significant change is observed in the performance of the gain compression inclusive model.

### 6.3 Sensitivity Analysis of DR-SCC

Sensitivity of the model to the hyperparameter is assessed (Figure 9). Sensitivity analysis to the ENR-Alpha hyperparameter, which increases the strength of the regularization with increasing value, demonstrates the extremes of the regularization (See Supplementary 6.4). When the ENR-Alpha is set very low *e*^*−*10^, reflecting low regularization, the readout does a better job of catching the week-to-week variations at the cost of including every oscillatory feature. Further work is needed to determine whether all SCC oscillations are important for depression severity and more nuanced measurements of behavior will be necessary.

**Figure 9:**
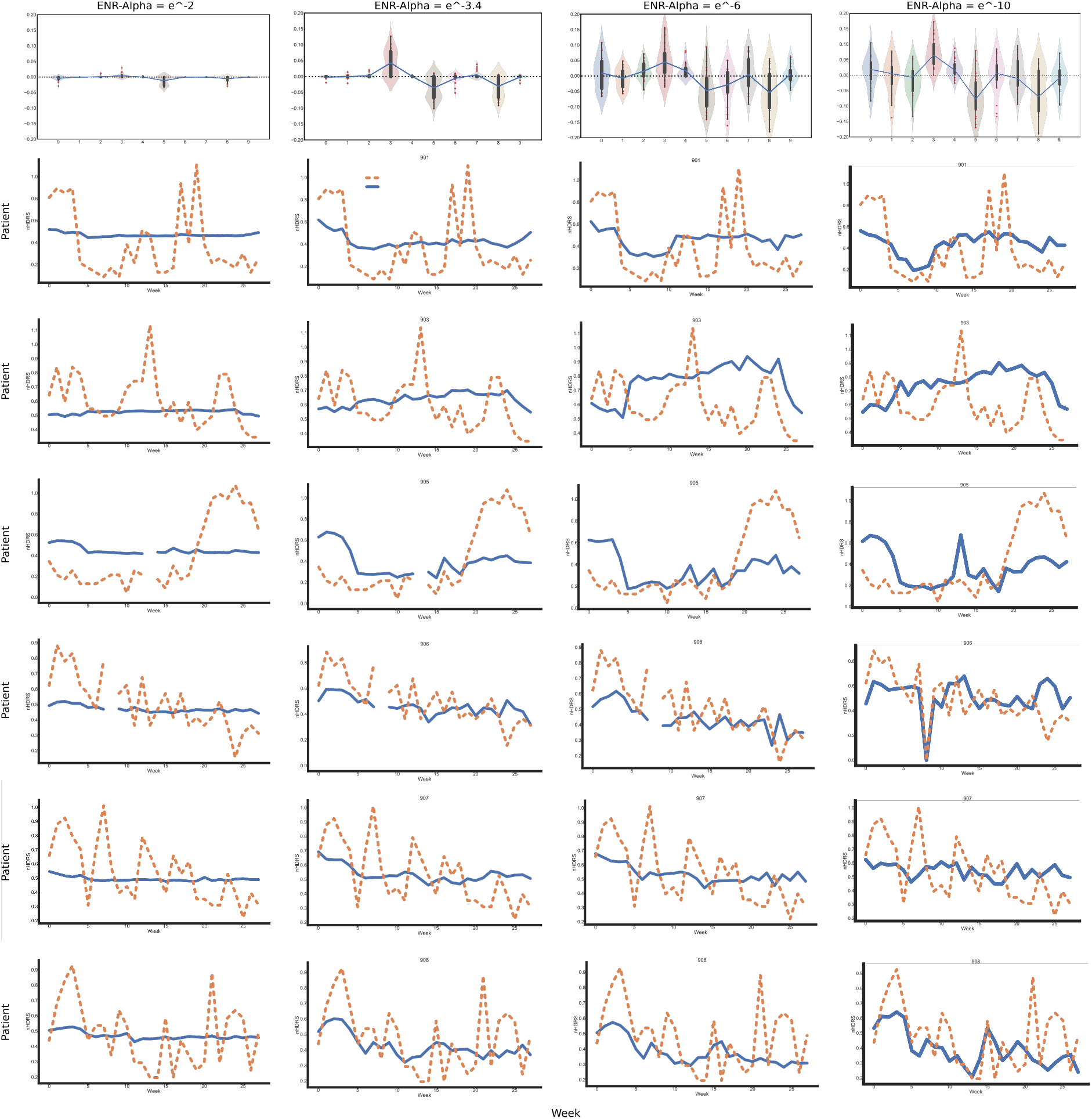
Sensitivity analysis for ENR-Alpha hyperparameter. Columns are ENR-Alpha = *e*^*−*2^, *e*^*−*3.4^, and *e*^*−*6^ respectively. Top row is the coefficients of the CV models and the final model (blue line). Next six rows are patient empirical nHDRS and SCC Readout superimposed. Final row is the scatterplot of predicted-vs-actual with associated statistics.

**Figure 10:**
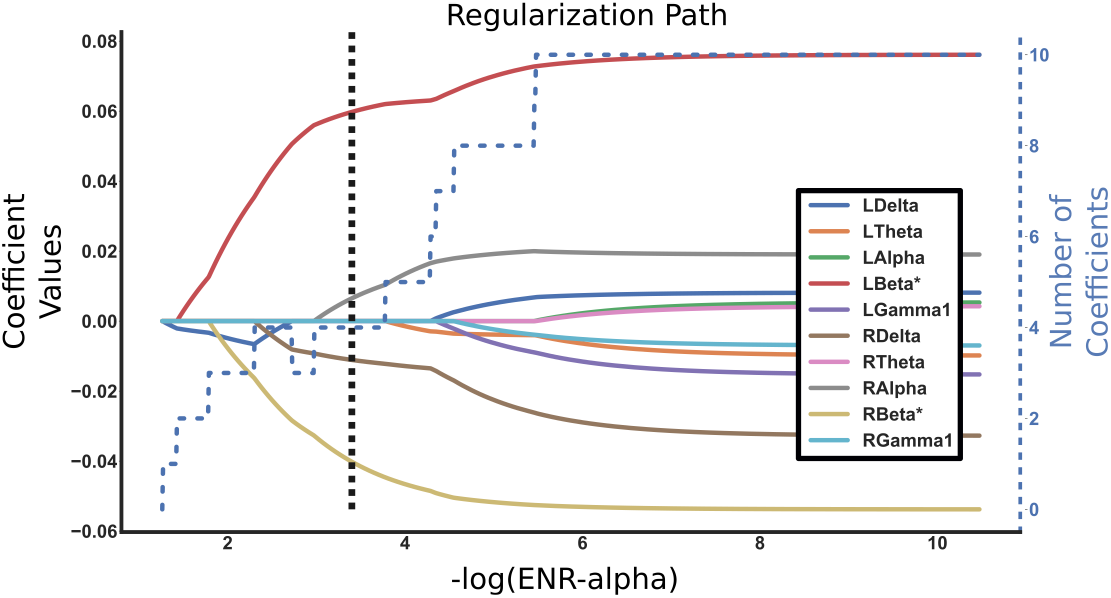
ENR Hyperparameter Selection Procedure. Regularization path depicts the coefficient values over varying levels of regularization. Number of non-zero coefficients is calculated (blue dotted line) along with regularization level. Final regularization level (black dotted line) is chosen at plateau of number of coefficients.

Visual inspection confirms broad fits to depression recovery, with DR-SCC missing transient weekly spikes in nHDRS that are of unclear origin. MDD is a slow-moving disorder and transient changes in emotion secondary to desired reactivity can manifest as large nHDRS spikes not requiring stimulation parameters adjusments [10].

### 6.4 Hyperparameter Selection

The hyperparameter for elastic net regularization (ENR) is chosen systematically after analysing the regularization path.

### 6.5 Synthetic Nulls

As a final confirmation that the model was statistically meaningful we used permutation testing and assessed our observed model performance compared to a set of synthetically constructed nulls. If the null hypothesis is true and there is no relationship between the SCC-state and the depression severity then the model should yield identical statistics on permuted testing-set data. 100 iterations of randomly shuffling the testing set nHDRS with respect to their weekly SCC state yielded a null distribution that the Predicted-vs-Actual slope and *R*^2^ would follow if there was no relationship (Figure11a,b). The statistics of the final model are then superimposed (black line Figure11a,b) and the number of null iterations that yielded a score higher than the actual model is used to compute the p-value. The model learned achieved statistical significance, further supporting the conclusion that our approach captures a significant relationship between SCC oscillations and depression severity that is disrupted by shuffling the nHDRS with respect to LFP.

**Figure 11:**
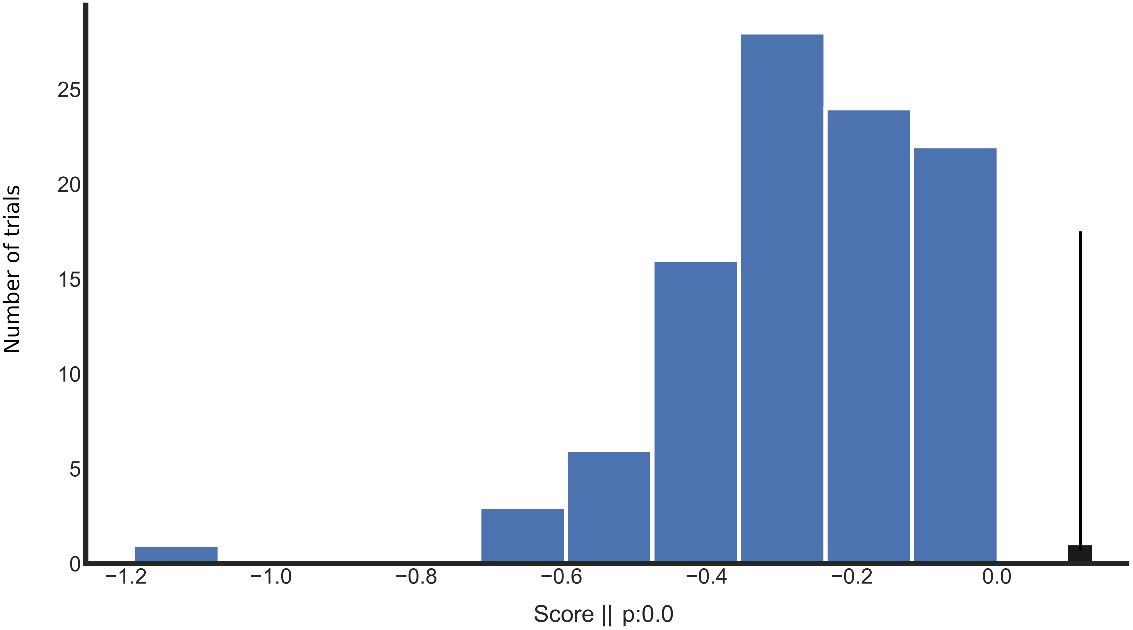
Null distribution of *R*^2^ over shuffled datasets. Synthetic null is constructed by shuffling weekly average SCC state against fixed nHDRS, destroying the putative relationship between the two variables. This is performed over 100 training iterations and the *R*^2^ in the testing set is calculated. DR-SCC *R*^2^ (black vertical line) is statistically significant.

